# RPA-PCR couple: an approach to expedite plant diagnostics and overcome PCR inhibitors

**DOI:** 10.1101/2020.03.02.969055

**Authors:** Mustafa Munawar, Frank Martin, Anna Toljamo, Harri Kokko, Elina Oksanen

## Abstract

Plant diseases are often diagnosed by the method of DNA extraction followed by PCR. DNA extraction from plant tissue can be a recalcitrant and lengthy process, and sometimes ends up with inhibitors that reduce PCR amplification efficiency. Here we present a unique approach, ‘RPA-PCR couple’, to exclude the DNA extraction step from the standard plant diagnostic process. The process crudely macerates plant tissue in water for a few minutes, and then transfers the macerate supernatant to a Recombinase Polymerase Amplification (RPA) reaction. Following an incubation of 20 minutes at 39 °C, the RPA reaction can be directly utilized in PCR amplification. In RPA-PCR couple, the RPA reaction is run at slower reaction kinetics to promotes amplification of long amplicons and the slower reaction kinetics are achieved by lowering RPA components concentrations. In this proof of concept study, we targeted *Phytophthora* intergenic mitochondrial spacer between *atp9* and *nad9* genes and the two common *Phytophthora* pathogens of strawberry: *P. fragariae* and *P. cactorum*. We presented coupling of RPA with real time TaqMan and SYBR Green PCR assays, and conventional PCR amplification aimed at Sanger sequencing. We found the RPA-PCR couple specific and capable of detecting as low as 10 fg of *Phytophthora* genomic DNA. Moreover, comparing RPA-PCR couple with the routine method of DNA extraction followed by PCR generated comparable results for the field samples. The idea of RPA-PCR couple to exclude DNA extraction may have vast application in different fields such as clinical diagnostics, food inspection and soil sciences.

## INTRODUCTION

A standard method for diagnosis of plant diseases is polymerase chain reaction (PCR). Traditionally, PCR requires prior extraction of DNA from infected tissue. Plant DNA extraction is not only a time-consuming process, but it also sometimes coextracts PCR inhibitors. To identify the presence of inhibitors in DNA samples, amplification of internal controls are often measured to ensure amplification efficiency is not affected. Although the internal controls spot false negatives, they also can increase cost and reduce amplification of the target amplicon when it is present in low concentrations. To combat reductions in amplification efficiency due to inhibitors, DNA extracts can be further purified, diluted, or have the DNA extraction repeated. While DNA dilution can reduce concentration of inhibitors, it also impacts the sensitivity of the assay by reducing the number of target DNA copies. For these reasons, DNA extraction from infected plant material may be a bottle neck in diagnosis of diseases (Martin et al., 2012).

An alternative DNA amplification method, Recombinase Polymerase Amplification (RPA), has been developed that addresses many of these limitations of DNA extraction (Piepenburg et al., 2006). RPA is an isothermal amplification where primers of 30-35 nucleotides length bind with recombinase enzyme, and these nucleoprotein complexes scan DNA to pair up with homologous region. This results in D-loop structure formation, where recombinase dissociates from primers and polymerase extend the primers. Bidirectional extension of primers by polymerase with strand displacement activity leads to exponential amplification of the target nucleic acid. Commercially available formulations for RPA assays can be obtained from TwistDx (United Kingdom), the patent holder of the technology, and Agdia (France). TwistDx, besides supplying reagents for RPA technology, aims at supporting the deployment of RPA technology in various fields including life sciences, diagnostics, microfluidics/lab-on-chip applications, agriculture, and defence.

RPA is reported to be tolerant of PCR inhibitors (Li et al., 2019) and can amplify DNA directly from crudely macerated plant tissue. Crude macerates have been prepared with different buffers. Ready-to-use buffers from Agdia, Inc. such as GEB (Kumar et al., 2018), GEB 2 (Miles et al., 2015; Li et al., 2017), GEB 4 (Zhang et al., 2014) and AMP1 (Kalyebi et al., 2015; Ghosh et al., 2018) have been utilized. Similarly, NaOH (Karakkat et al., 2018; Qian et al., 2018), standard ELISA grinding buffer (Miles et al., 2015), and TE buffer (Ahmed et al. 2018) have also been used for maceration. Besides maceration, extraction buffers have been employed to release nucleic acid from leaf disks (Si Ammour et al., 2017; Silva et al., 2018). Moreover, squeezed sap has been directly utilized for RPA (Wambua et al., 2017).

Besides tolerances to inhibitors, RPA is also rapid and sensitive, and it requires simple instrumentation. For these reasons, RPA has gained attention from plant pathologists, but users may encounter some challenges when designing amplification primers with the necessary specificity. Although many researchers have reported satisfactory specificity of their RPA assays, some have also reported cross reactivity (Patel et al., 2016; Yang et al., 2017; Moore and Jaykus, 2017). RPA assays have also been reported tolerating 5, 7 and even 9 mismatches with target sites (El Wahed et al., 2013; Boyle et al., 2013; Daher et al., 2015). The study reporting tolerance up to 9 mismatches originally investigated the effect of different mismatches on RPA specificity and recommended selection of a target marker with at least 36% sequence difference with the closely related species while designing RPA assays (Daher et al., 2015). RPA primers, in contrast to PCR, do not depend on melting temperatures to anneal with target sites. Amplification at low temperature ranging from 25-42 °C may enable RPA primers to make dimers and even bind to matching non-target sites, and thereby generating nonspecific by-products that can be visible on an agarose gel. The nonspecific products compete with the target amplification for the limited availability of energy available in the RPA reaction. This background amplification may also complicate amplification of longer templates; sensitive amplification generally requires small amplicons, ideally of 100-200 bp. Longer amplicons require lengthier extension time and thereby target amplification may be outcompeted by the background amplification. For longer amplicons, TwistDx manuals recommend slower reaction kinetics through lowering incubation temperature and concentration of magnesium acetate and primers. In our experience, lowering concentration of magnesium acetate and primers is beneficial for longer amplicons, but detection of low amount of longer targeted amplicons (∼ 500 bp) is still not possible. It should be noted that background amplification is less problematic for the RPA chemistries designed for inclusion of a probe-based detection system, such as TwistAmp^®^ exo kit.

While both PCR and RPA have their disadvantages, a unique combination of them can result in improved detection capabilities. Coupling slower reaction kinetics RPA with PCR can exclude the need for time consuming and problematic DNA extractions. The method is also expected to overcome the RPA shortcomings of non-specificity (background amplification), short amplicon size (100-200 bp optimal) and poor resolution for quantification. In the coupled amplification, RPA first multiplies the template in the first-round amplification directly from plant macerate. The amplicons from the first round are then directly amplified by PCR. Slower RPA reaction kinetics promote generation of longer amplicons. Similarly, the slower rate of RPA amplification generates less background noise and target amplicons which can be directly utilized for PCR (without dilution). The coupled nested PCR are run with normal chemistry.

In this this proof-of-concept study, we will validate ‘RPA-PCR couple’ as a simple, rapid, sensitive and specific approach to directly amplify target DNA from plant crude macerates. The study will target *Phytophthora* intergenic mitochondrial marker *atp9-nad9* and the concept will be applied to two *Phytophthora* species causing devastating damages to strawberry. The pathogens will include *Phytophthora fragariae*, causative agent of red stele of strawberry roots, and *Phytophthora cactorum*, agent of strawberry crown rot. Our first round RPA assay will be *Phytophthora* genus-specific and it will amplify the intergenic spacer between *atp9* and *nad9* gene at slower reaction kinetics. Then the RPA reaction will be coupled either with species-specific real-time PCR assays, or genus-specific conventional PCR aimed at Sanger sequencing. The species-specific assays include SYBR Green and TaqMan assays for *P. fragariae* and *P. cactorum*, and their specificity originate from the intergenic spacer between *atp9-nad9*.

## MATERIAL AND METHODS

### RPA Primers Screening

Miles et al. (2015; Supplemental File 2) provided 274 partial *atp9-nad9* sequences of approximately one hundred *Phytophthora* species. The sequences were aligned by Geneious 8.1.9 (Geneious, New Zealand) to identify the *atp9* gene, intergenic spacer, and *nad9* gene sequences. Ten forward RPA primers were picked from *atp9* region and ten reverse primers from *nad9* region. Primers were screened to find the optimal RPA primer pair generating the maximum amount of the target band (∼ 500 bp). DNA extracted from *P. fragariae* hyphae was utilized as template and the DNA volume was kept at 1 µl per reaction. DNA extraction from *Phytophthora* hyphae was previously described (Munawar et al., 2019). The concentration of the hyphal DNA was 100 pg/µl. All DNA concentrations in this study were quantified through Qubit 2.0 Fluorometer, utilizing Qubit dsDNA HS Assay Kit (Thermo Scientific, Unites States).

In primers screening, RPA reactions were run at normal reaction kinetics utilizing the ‘TwistAmp^®^ Liquid Basic’ kit (TwistDx Inc., UK.) and following manufacturer recommendations. RPA incubation was done at 39 °C for 25 min with no agitation and using a PTC-0200 DNA Engine Cycler (BIO-RAD, United States) with no lid heating. Following incubation, reactions were purified with a GeneJET PCR Purification kit (Thermo Scientific, United States) and results were examined by the end point determination method of gel electrophoresis. The forward RPA primer ‘ATP9F-M9’ and reverse RPA primer ‘Phy_Gen_R20’ (Table 1) were found to be the optimum RPA primer pair (data not shown).

**Table 1:**
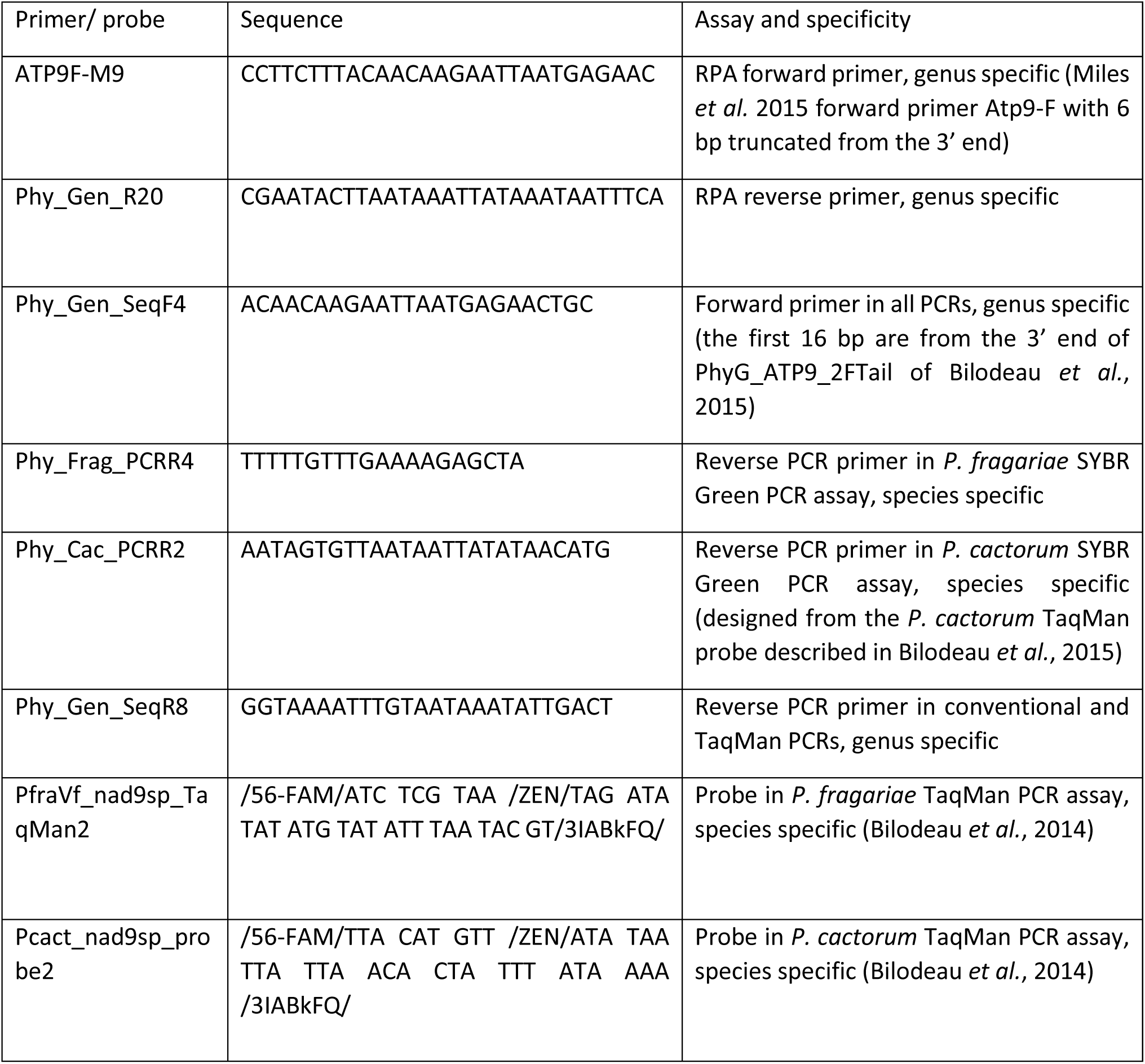
Primers and probe sequences and role.

### Real Time SYBR Green PCR Assays

All PCR reactions in this study were performed with 500 nM primer concentration and an amplification volume of 25 µl. PCR primers were designed through Geneious 8.1.9 where default GC % and Tm were often lowered. All quantitative PCR amplifications were run with Mx3000P QPCR System (Agilent, Germany). All primers and probes were ordered from Integrated DNA Technologies, Inc (Belgium). Primers and probe sequences are provided in Table 1, with their locations in the *atp9-nad9* region presented in Figure 1.

**Figure 1:**
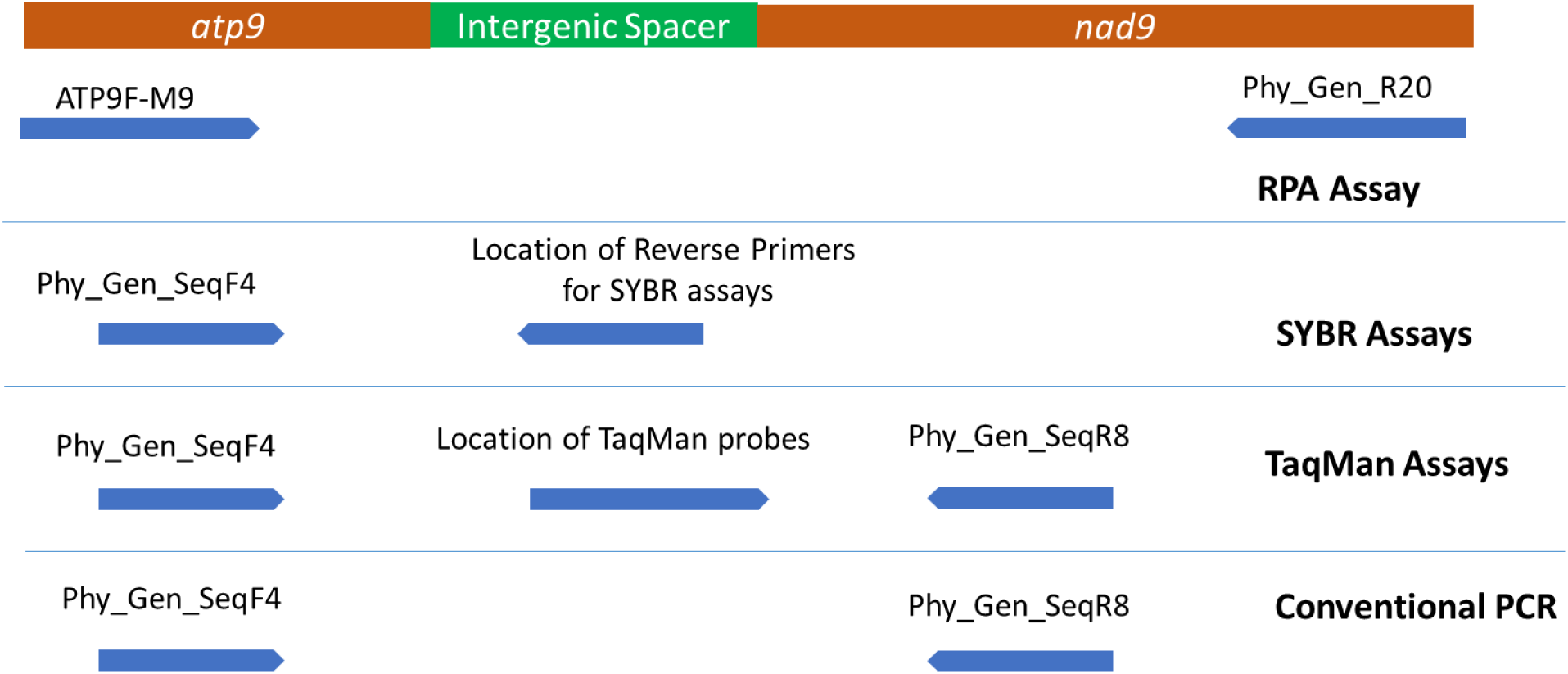
Location of primers in RPA, SYBR Green PCR, TaqMan PCR and conventional PCR assay.

In species specific SYBR Green PCR assays, reverse primers were designed from the intergenic spacer between the *atp9* and *nad9* genes, and forward primers from the conserved region of *atp9*. In the *P. fragariae* assay, the forward primer ‘Phy_Gen_SeqF4’ (first 16 bp are from the 3’ end of PhyG_ATP_3F of Bilodeau et al., 2014) and reverse primer ‘Phy_Frag_PCRR4’ were utilized. The amplification program included 10 min pre-denaturation at 95 °C, and 40 amplification cycles, where each cycle has 10 sec denaturation at 95 °C, 20 sec annealing at 58 °C and extension of 20 sec at 67 °C. The *P. cactorum* assay was done with the forward primer ‘Phy_Gen_SeqF4’ and reverse primer ‘Phy_Cac_PCRR2’. Amplification conditions were similar to *P. fragariae* with the exception of a 55 °C annealing temperature, and extension of 30 sec at 60 °C. All SYBR Green assays were completed with LightCycler^®^ 480 SYBR Green I Master (Roche, Switzerland). Melt curve analysis was always included in SYBR Green amplifications.

### Real Time TaqMan Assays

Bilodeau et al. (2014) designed and validated species specific TaqMan assays for several *Phytophthora* species including those used in this experimentation. In these assays, probes were placed in the intergenic spacer between the *atp9* and *nad9* genes, while forward and reverse primers were designed from the flanking genus conserved regions. The probes for *P. fragariae*, ‘PfraVf_nad9sp_TaqMan2’, and for *P. cactorum*, ‘Pcact_nad9sp_probe2’ were ordered as ZEN Double-Quenched Probes to improve signal to background discrimination. The forward primer ‘PhyG_ATP9_2FTail’ and reverse primer, ‘PhyG-R6_Tail’ were replaced with forward primer ‘Phy_Gen_SeqF4’ and reverse primer ‘Phy_Gen_SeqR8’ as the latter improved sensitivities. In the species specific TaqMan PCR assays, probe concentration was kept at 100 nM and Luminaris Probe q PCR Master Mix (Thermo Scientific) was used. The PCR program for the *P. fragariae* assay started with a 2 min incubation at 50 °C to let Uracil-DNA Glycosylase (UDG) remove possible carryover contamination. Then a 10 min incubation at 95 °C to activate Hot Start Taq DNA Polymerase followed by 45 amplification cycles. Each cycle consisted of a denaturation step of 15 sec at 95 °C, and an annealing/ extension step of 90 sec at 57 °C. The amplification program for the *P. cactorum* assay was the same but with an annealing/ extension step at 55 °C.

### Conventional PCR for Sequencing

Conventional PCR amplification of the *Phytophthora* genus specific amplicon was done to generate template for Sanger sequencing. The amplification utilized the previously mentioned ‘Phy_Gen_SeqF4’ and ‘Phy_Gen_SeqR8’ primers. LightCycler^®^ 480 SYBR Green I Master was utilized as master mix due to its exceptional capability to amplify even after 50 cycles. The fluorescence signals were not considered, rather gel electrophoresis was utilized to observe amplification of the target ∼ 300 bp bands. The amplification program started with a 10 min pre-incubation at 95 °C and followed 50 amplification cycles of 10 sec at 95 °C, 20 sec at 55 °C, and 30 sec at 65 °C. PCR reactions with visible target bands were purified with GeneJET PCR Purification and sent for Sanger sequencing to GATC Biotech, Germany.

### Coupling RPA with PCR

For the RPA-PCR couple method, ‘TwistAmp^®^ Liquid Basic’ kit was used to prepare RPA reaction mix for amplifying a target of around 500 bp under conditions of slower reaction kinetics. A pre-master mix was prepared in a 1.5 ml tube. The pre-master mix 1X recipe included 25 µl of 2x Reaction Buffer, 2 µl 10x Basic E-mix, 1 µl of 40 mM dNTPs (total), 1.5 µl forward primer ‘ATP9F-M9’ (10 µM), 1.5 µl of reverse primer ‘Phy_Gen_R20’ (10 µM), and 15.21 µl water. The pre-master mix was vortexed and spun briefly, and the 20x Core Reaction Mix was added to the lid and the tube was inverted 10 times to prepare final master mix. For 1X recipe of the master mix, 20x Core Reaction Mix volume was kept at 1 µl. The 20x Core Reaction Mix was also warmed to room temperature and pipette mixed before use. Then 47.21 µl of the master mix was added to individual amplification tubes followed by 1 µl macerate/template and 1.79 µl of 280mM MgOAc that were added to the inside of the lid strip. Reactions were started by inverting the tubes to mix and the amplification mixture concentrated at the bottom of the tube prior to immediately incubating the tubes at 39 °C for 20 min, with an agitation step after four minutes. Following 20-minute incubation, reaction enzymes were denatured by heating at 85 °C for 1 min to terminate amplification and cooling to 4 °C. The RPA incubation was done in a PTC-0200 DNA Engine Cycler with no lid heating. Once the RPA reaction was completed, 1 µl reaction was directly utilized for PCR or frozen at -20 °C for future use. In RPA-PCR couple, the number of PCR amplification cycles were reduced. The coupled species specific SYBR Green and TaqMan amplifications were done with 30 thermal cycles, while conventional PCR genus-specific amplification was done with 40 cycles.

For estimation of sensitivity of the RPA-PCR couple method, slower reaction kinetics RPA reactions were done with 1 µl of *P. fragariae* or *P. cactorum* hyphal DNA dilutions at concentrations ranging from 100 pg/µl to 0.1 fg/µl and coupled with related PCR assays. The RPA reactions with *P. fragariae* DNA dilutions were spiked with macerate of healthy rootlet tips, while reactions with *P. cactorum* DNA were spiked with macerate of healthy crown tissue. Macerate preparation was previously described (Munawar et al., 2019) and added in amounts that would be used for assaying plant samples (1 µl).

### Specificity

NCBI Primer-BLAST was customized to analyse specificity of the species level SYBR Green PCR assays designed in this study. In the analysis, the partial *atp9-nad9* sequences of approximately one hundred *Phytophthora* species provided in Miles et al. (2015; Supplemental File 2), and complete *atp9* gene sequences from 28 *Phytophthora* species received from United States Department of Agriculture, Agricultural Research Service, Salinas, USA were utilized. In laboratory evaluations of specificity, DNA extracts of *Phytophthora fragariae, Phytophthora cactorum, Phytophthora taxon raspberry, Phytophthora megasperma, Phytophthora rosacearum, Phytophthora ramorum, Phytophthora plurivora, Phytophthora pini, Phytophthora cambivora, Phytophthora cinnamomi, Pythium sylvaticum, Botrytis cinerea, Colletotrichum acuta*tum, *Mucor hiemalis, Fusarium avenaceum*, and *Fusarium proliferatum* were utilized. Details of the taxa included in laboratory evaluations of specificity are provided in Table S1 in supportive information. DNA extracts concentrations ranged between 50 pg/ µl and 100 pg/ µl and 1µl of each DNA extract was added in individual reaction tubes. We first analysed all the PCR assays individually for possible cross reactivity with DNA from non-target taxa. Then we analysed RPA-PCR couple specificity by adding DNA from each taxa individually for RPA amplification and coupling those RPA reactions with all formats of PCR assays.

### Field Validation

For field validation, 22 fine rootlet samples and 22 crown samples were collected from plants investigated for red stele and crown rot disease in the summer 2018. The plants samples were from 25 different problematic fields of Northern Savo region of Finland. Each rootlet sample was collected from three plants from a field (pooling of finest root tips). In contrast, crown samples were not pooled, rather each crown sample was collected from individual plants. The crown samples always included rot if it was observed. Tissue samples weighing from 25-100 mg were collected in round bottom 2 ml Eppendorf tube and macerated in autoclaved distilled water with a plastic pestle. Rootlet macerates were prepared as 1:10 (w/v) tissue weight by volume of water, while crowns macerates were 1:20 (w/v). To facilitate maceration, initially only 100 µl water was added and following maceration, more water was supplemented to achieve final desired dilution. Once macerates were ready, they were vortex and left for particulate settlement for around one minute. The maceration process is explicitly explained in Munawar et al. (2019). From tissue macerates, 1 µl of supernatant was transferred to RPA amplifications done with slower reaction kinetics. Once the RPA amplification was completed, RPA reactions were coupled with the four species-specific real-time PCR assays and one conventional genus-specific PCR assay. The target bands from conventional PCR were also Sanger sequenced to confirm identification of the taxa present.

To validate RPA-PCR couple results and compared them with standard diagnostic procedure of DNA extraction-PCR, the macerates prepared were frozen at – 80 °C. Later the samples were freeze dried, and DNA extracted as reported by Till et al. (2015), directly starting from the step fifth of lysis buffer addition. In DNA extraction, we utilized 100 µl of liquid silica stock per tube, guanidine thiocyanate as DNA binding buffer and LB2 as lysis buffer. Following DNA extraction, DNA volumes of 0.25 µl, 0.5 µl and 1 µl from each sample were tested with the four species-specific and one conventional PCR amplification. The products from conventional PCR were also Sanger sequenced. All DNA samples which showed no amplification were tested for presence of PCR inhibitors or DNA extraction failure through a separate control PCR amplification targeting plant DNA. The control PCR was a SYBR Green PCR assay targeting plant sequences that included the FMP12b and FMP13b primers of Tooley et al. (2006), master mix LightCycler^®^ 480 SYBR Green I Master, and 0.25 µl DNA samples. The amplification program started with 10 minutes incubation at 95 °C followed by 35 amplification cycles of 95 °C for 10 seconds, 55 °C for 20 seconds, and 72 °C for 20 seconds.

## RESULTS

### Sensitivity and Specificity of PCR assays

Both species specific SYBR Green assays detected as low as 1 fg of target genomic DNA (eight replicates) with linearity down to 10 fg. The melt curves produced double peaks as predicted by uMelt (https://www.dna.utah.edu/umelt/umelt.html). Amplification efficiency of the *P. fragariae* assay was 98.76 % and the *P. cactorum* assay was 94.50 %. Like the SYBR Green assays, both species specific TaqMan assays amplified as low as 1 fg of target genomic DNA (eight replicates) with linearity down to 10 fg. Amplification efficiency of the *P. fragariae* assay was 90.10 % and the *P. cactorum* assay was 82.91 %. The conventional PCR genus-specific assay produced visible target band of ∼ 300 bp for as low as 1 fg *Phytophthora* hyphal DNA (two replicates).

Primer-BLAST predicted cross reactivity of the *P. fragariae* SYBR Green assay with only *Phytophthora alni, Phytophthora cambivora*, and *Phytophthora rubi*. Similarly, Primer-BLAST foretold cross reactivity of the *P. cactorum* SYBR Green PCR assay with *Phytophthora pseudotsugae, Phytophthora hedraiandra*, and *Phytophthora idaei*. In laboratory analysis of specificity, the species-specific TaqMan and SYBR Green assays showed no cross reactivity for the taxa tested with the exception of the SYBR Green PCR assay for *P. fragariae*, which cross reacted with *P. cambivora* as predicted by Primer-BLAST. Similarly, the conventional genus-specific PCR amplified all *Phytophthora* species and showed no cross reactivity with any of the non-*Phytophthora* species DNA.

### Sensitivity and Specificity of RPA-PCR couple

For sensitivity evaluation, slower reaction kinetics RPA reaction were done separately for 1/10 serial dilutions of *P. fragariae* and *P. cactorum* hyphal DNA. The RPA reactions were also spiked with healthy rootlets and crown macerate, respectively. Following amplification, aliquots of the RPA reactions were transferred to real time TaqMan and SYBR Green PCR assays. The RPA-PCR couple consistently amplified target *Phytophthora* hyphal DNA down to 10 fg starting concentration (six replicates). Figure 2 represents standard curves when slower kinetics RPA reactions were done with 1/10 serial dilution of *P. fragariae* DNA and healthy rootlets macerates, and the RPA reactions were coupled with *P. fragariae* specific SYBR Green PCR assay. Similarly, Figure 3 represents a similar set up of RPA-PCR coupled where RPA reaction were used with dilutions of *P. cactorum* DNA and spiked with healthy crown macerate, and later coupled with *P. cactorum* specific TaqMan PCR assay.

**Figure 2:**
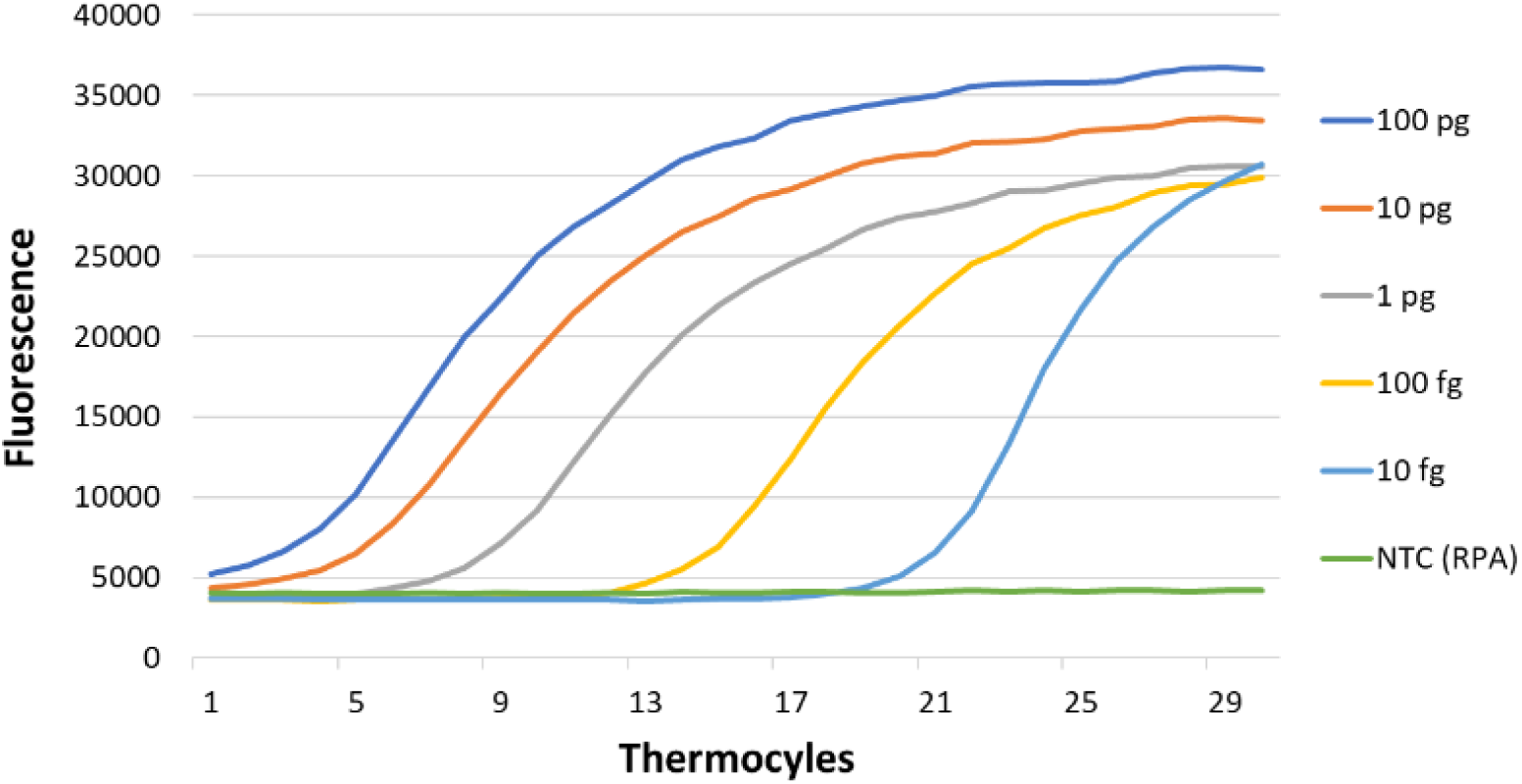
Standard curve of RPA-PCR couple when template was *P. fragariae* DNA and the coupled PCR is *P. fragariae* specific SYBR Green PCR assay. The curves represent 1:10 serial dilutions of DNA from 100 pg to 10 fg used in the RPA amplification.

**Figure 3:**
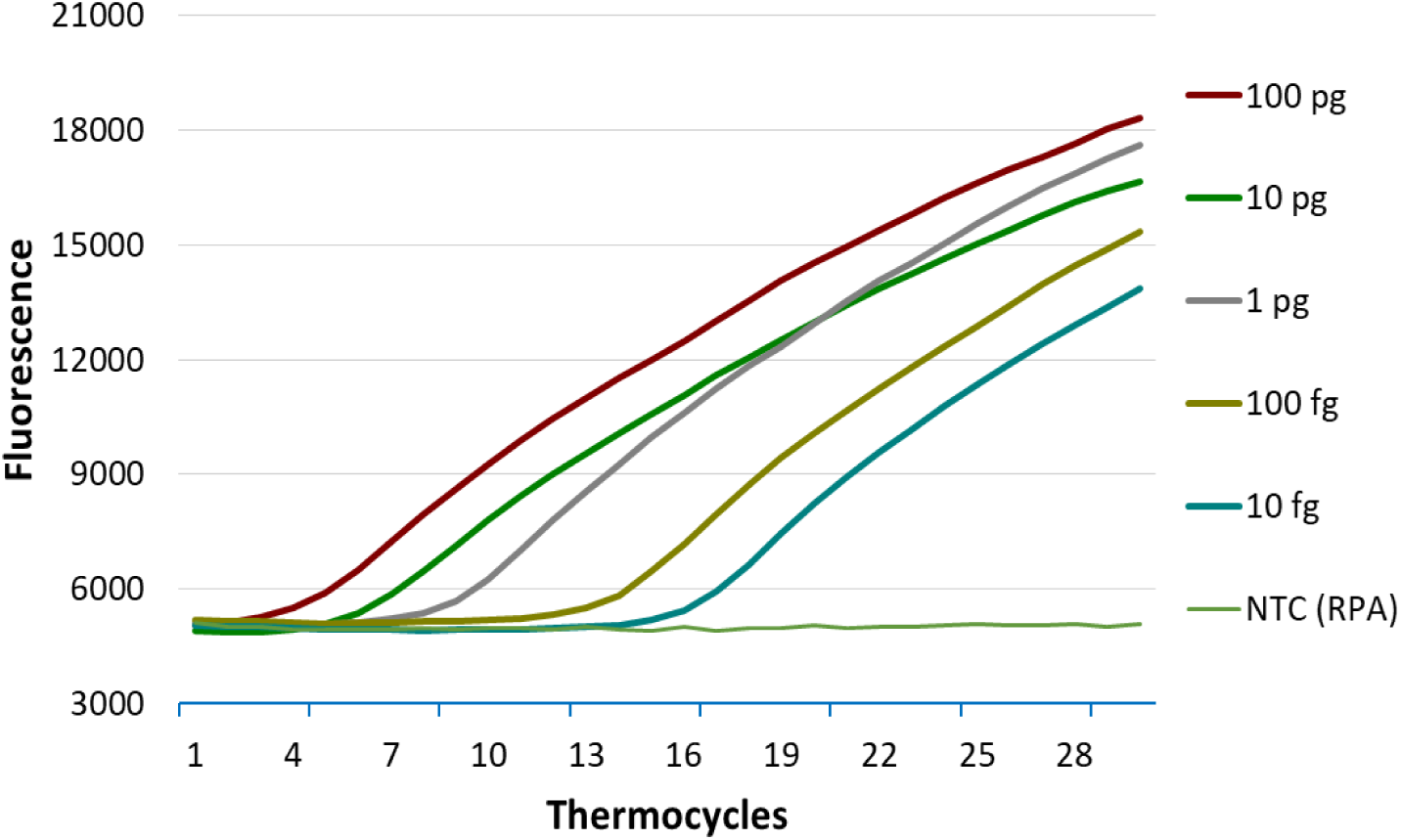
Standard curve of RPA-PCR couple when template was *P. cactorum* DNA and the coupled PCR is *P. cactorum* specific TaqMan PCR assay. The curves represent 1:10 serial dilutions of DNA from 100 pg to 10 fg used in the RPA amplification.

In laboratory specificity analysis of RPA-PCR couple, RPA amplification was done with DNA from non-target taxa and later coupled with the four real time PCR and one conventional PCR assays. We observed that only the RPA reaction with *P. cactorum* DNA was detected with the coupled *P. cactorum* specific real time TaqMan and SYBR Green PCR assays. Similarly, only the RPA reaction with *P. fragariae* was detected with the coupled *P. fragariae* specific real time TaqMan PCR assay, but the coupled *P. fragariae* real time SYBR Green also detected *P. cambivora* besides *P. fragariae* as expected. Coupling the RPA reactions with genus specific conventional PCR amplified all the *Phytophthora* species with no amplification observed with other taxa. Sequencing the *Phytophthora* amplicons confirmed their species identification.

### Field Validation

For field validation, we compared RPA-PCR couple method with the method of DNA extraction followed by PCR. We observed some difference in the results of the two methods for the field samples, but all the five versions of PCR generated consistent results within each method. The five PCR assays were *P. fragariae* specific TaqMan PCR, *P. fragariae* specific SYBR Green PCR, *P. cactorum* specific TaqMan PCR, *P. cactorum* specific SYBR Green PCR, and *Phytophthora* genus specific conventional PCR (Sanger sequencing).

In the process of validation and comparison, we first did RPA-PCR couple from macerates (one day process) and later extracted DNA from the macerates to perform PCR assays (two days process). All the DNA extracts which gave no amplification with the five target PCR assays were further tested with a control PCR to exclude possibility of inhibitors or extraction failure. For all those samples, control PCR generated Ct value between 22 to 28, except one crown DNA which had a Ct around 33. The crown sample with the high Ct was excluded from validation process. Regarding PCR from DNA extraction, all crown DNA samples amplified with 0.5 µl and 1 µl volume, while rootlets DNA samples had more inhibitors and volumes of 0.25 µl and 0.5 µl were mostly successful.

In the comparison of RPA-PCR couple with the method of DNA extraction followed by PCR, the later produced more positives results. Through RPA-PCR couple, among the 22 rootlet samples investigated, nine were found infected with *P. fragariae*, four with *P. cactorum*, and one co-infected with both *P. fragariae* and *P. cactorum*. The coinfection was detectable through species specific PCR assays, but Sanger sequencing was not successful for that sample. Similarly, analysis of 21 crown macerates with RPA-PCR couple revealed that eight were infected with *P. cactorum*, while 13 crown macerates gave no amplification. Regarding results of PCRs from DNA extractions, all positives samples from RPA-PCR couple were also found positive. PCR assays from DNA extractions detected three additional weak or late positive samples: one rootlet sample infected with *P. cactorum* and two crown samples contaminated with *P. fragariae*. The Sanger sequencing of the co-infected rootlet samples didn’t fail, rather generated good reads from *P. fragariae atp9-nad9* marker (possibly due to bias of PCR). All the Sanger sequences of *P. fragariae* were identical to JF771842 (GenBank accession number) and all sequences of *P. cactorum* were identical to MH094138 (GenBank accession number).

## DISCUSSION

In this study, we were able to demonstrate that RPA-PCR couple is a time-saving procedure to amplify target nucleic acid directly from simple plant macerates. Through RPA-PCR couple, one can specifically amplify target nucleic acid of sufficient length without the need of extracting DNA from plant tissue. RPA-PCR couple is about crudely macerating plant tissue with water in a few minutes and then doing a 20-minute RPA (slower reaction kinetics) from macerates. Following RPA amplification these reactions can be directly utilized for PCR amplification, while the PCR can be a conventional, TaqMan or SYBR Green real time PCR assay. In RPA-PCR couple, the RPA amplifies target nucleic acid directly from plant macerate. Similarly, when RPA is run at slower reaction kinetics, it amplifies long target amplicons such as 500 bp. Although slower reaction kinetics of RPA promotes amplification of long target sequences, it also generates limited copies of the amplicon. These amplicons are further replicated by the PCR and the replication is specific and exponential. RPA-PCR couple will be able to amplify both DNA and RNA as the RPA technology can amplify both types of nucleic acids. Regarding cost of RPA-PCR couple, the cost of RPA (TwistAmp^®^ Liquid Basic) kit is comparable with standard plant DNA kits. Although the idea of RPA-PCR couple was conceived for plants, it may work for other samples encountering the PCR inhibitors problem when DNA extracted, such as food or faecal samples (Schrader *et al*., 2012).

We initially aimed at direct amplification of *Phytophthora atp9-nad9* marker from infected plant tissue through RPA to proceed with Sanger sequencing, but we faced problems. To succeed with Sanger sequencing, we required to gel cut the target band and often the concentration of the target amplicon was not enough. So, we switched to coupling the RPA with PCR. The concept of coupling RPA with PCR was earlier presented by Miles *et al.* (2015) where standard RPA reaction (normal reaction kinetics) was coupled with PCR to sequence and validate some RPA results. We observed coupling of RPA reactions (lyophilized ‘TwistAmp Basic’ kit) with PCR for our long target of around 500 bp length were not successful for the low target concentrations. Moreover, excessive amplification in RPA reaction, either target or non-specific, required dilution of the RPA reaction before PCR. Following TwistDx recommendations for long amplicons, we observed lowering magnesium acetate and primers favoured amplification of long target to some extent, but the issue was resolved through liquid RPA basic kit (TwistAmp^®^ Liquid Basic) which provides flexibility to adjust RPA reaction components. By slowing RPA reaction kinetics in the RPA-PCR couple, we succeeded amplifying our long target at femto levels. Although the slower reaction kinetics RPA generates a limited amount of the target, the low quantity of amplicons is not problematic as the coupled PCR exponentially replicates those amplicons. To achieve slower reaction kinetics in TwistAmp^®^ Liquid Basic kit, concentrations of primers, MgOAc, ATPs (10X basic E-Mix), building blocks (dNTPs) and related enzymes (20X core reaction mix) were lowered. TwistAmp^®^ Liquid Basic is provided with components of 2X reaction buffer, 10X Basic E-Mix, 20X core reaction mix, and 280mM MgOAc. The 2X reaction buffer comprises of high salt buffer, Tris, Potassium Acetate, PEG, and dH_2_O. Similarly, the 10X Basic E-Mix contains phosphocreatine, ATP, Creatine Kinase (CK), high salt buffer, and dH_2_O. The 20X core reaction mix comprises GP32, UvsX, UvsY, and Polymerase (communication with TwistDx).

RPA-PCR couple is more than a combination of two nucleic acid amplification technologies. RPA-PCR couple is time saving. Running RPA prior to PCR provides the possibility to amplify target from simple macerates and thereby excludes the need for the time-consuming procedures of plant DNA extraction or PCR inhibitor removal. In RPA-PCR couple, a 20-minute RPA is coupled with PCR, and the coupled PCR also run for fewer cycles (lesser time). In our observation, the slower reaction kinetics of RPA provides a better resolution for quantification through real-time quantitative PCR assays (compared to standard RPA assays such as TwistAmp^®^ exo assay). The slower reaction kinetics of RPA also facilitates amplification of long targets (over 500 bp) with improved sensitivity. The slower reaction kinetics excludes the need for diluting RPA reaction prior to doing PCR. In RPA-PCR couple, the background noise generated in RPA reaction is also less problematic; limited background noise is generated due to the slower reaction kinetics and the coupled PCR takes the role of target amplification in the second cycle of amplification. While RPA and other isothermal nucleic acid amplification technologies often suffer from background amplification problems, their utilization for initial and limited amplification of target nucleic acid to replace the DNA extraction step provides new insight.

In our RPA-PCR couple optimization, we first screened RPA primers at normal RPA reaction kinetics, then optimized PCR assays and finally tested the coupling. For RPA primers screening, we had to run RPA at normal reaction kinetics as the slower reaction kinetics generates limited quantity of amplicons and low quantity is not observable through gel electrophoresis. Moreover, we utilized high concentration of initial template (100 pg/µl *Phytophthora* hyphal DNA) in RPA primer screening as our template was ∼ 500 bp long, while RPA normal reaction kinetics favour amplification of shorter targets. For RPA primer screening, we employed the conventional method of gel electrophoresis which requires RPA reactions to be cleaned prior gel electrophoresis. We suggest that one can also adopt a reverse approach for optimizing RPA-PCR couple, meaning that first optimizing a PCR assay and later utilizing the PCR assay to compare the slower reaction kinetics RPA reactions added with different combination of RPA primers. Screening RPA primers through the coupled real time PCR will exclude the need of RPA reaction clean up and will provide better data to compare.

RPA-PCR couple standard curves were not as steep and equally spaced as of real time PCR assays. The slight decrease in RPA-PCR couple curves steepness possibly indicates inhibition of PCR by RPA reagents. Moreover, as RPA is run first in RPA-PCR couple, the PCR curves are more representative of the RPA reactions done from 1/10 serial dilution of *Phytophthora* hyphal DNA and therefore curves are not equally spaced. In laboratory specificity analysis of RPA-PCR couple, DNA of the different taxa was diluted to a range between 50 pg/ µl and 100 pg/ µl. Addition of DNA more than 100 pg in an RPA reaction tends to give false amplification in the coupled PCR assays, especially SYBR Green assays, but any false positive amplifications in SYBR Green PCR assay can be identified through melt curve analysis. Moreover, such high DNA concentrations are not generally present in plant crude macerates. Regarding field validation and comparison of RPA-PCR couple with the routine method of DNA extraction followed PCR, RPA-PCR couple missed few weak positive samples. This probably happened due to lesser sensitivity of RPA-PCR couple. RPA-PCR couple consistently detected as low as 10 fg *Phytophthora* hyphal DNA, while PCR assays were detecting as low as 1 fg of the DNA. Moreover, RPA targeted longer amplicon of around 500 bp, while PCRs amplified shorter targets of around 200-300 bp.

Optimizing PCR assays designed in this project from the low GC *atp9-nad9* marker was also challenging, especially in the case of *P. cactorum*. In the *P. cactorum* SYBR Green assay, the extension temperature was lowered to 60 °C and additional 10 sec were provided to assist completion of extension step. In *P. cactorum* TaqMan assay, the ‘Pcact_nad9sp_probe2’ has a GC content of only 11.1 % and Tm of 53.8 °C. Even after reducing annealing/ extension temperature to 55 °C, the amplification curves were more linear and missing the plateau phase. Ideally the TaqMan probe for *P. cactorum* probe used in this study should be re-designed with addition of MGB (minor groove binder) to raise Tm to adequate level. Otherwise the TaqMan probe should be further elongated to raise Tm. Regarding the *Phytophthora* genus-specific conventional PCR, extension temperature was lowered to 65 °C. An extension temperature of 68 °C worked for *P. fragariae* but failed for *P. cactorum*.

Applying the method to universal identification markers can expedite species identification. In the current study, utilization of the unique intergenic *atp9-nad9* marker provides a rapid universal *Phytophthora* genus and species-specific assay with potential to identify almost all *Phytophthora* species (Bilodeau *et al.*, 2012, Miles *et al.*, 2017). The universal assay is about crudely macerating *Phytophthora* infected plant tissue in water and transferring the macerate supernatant into RPA reaction tuned for slower reaction rate. The RPA reaction added with forward primer ‘ATP9F-M9’ and reverse primer ‘Phy_Gen_R20’ amplifies the *atp9-nad9* marker. The RPA reaction is transferred to conventional PCR assay (‘Phy_Gen_SeqF4’ and ‘Phy_Gen_SeqR8’ primers) which in turn exponentially amplifies the target marker. Sanger sequencing the cleaned PCR product identified the *Phytophthora* species. It is notable that this universal assay will not be applicable to *Phytophthora bisheria* or *Phytophthora frigida* due to the possible unusual gene order of *atp9* and *nad9* genes (Bilodeau *et al.*, 2014).

Although RPA generate limited product in the RPA-PCR couple, opening RPA tubes to transfer reactions into PCR pose risk of contamination. RPA strip opening and reaction transfer should be carefully accomplished in some suitable hood of post-PCR laboratory. Similarly, filter tips or positive displacement tips should be utilized to avoid contaminations. Good molecular laboratory practice of physically separating pre and post amplification laboratories, preparing macerates and reactions reagents in pre-amplification laboratory, and including No Template Control (NTC) guarantee quality of results. RPA reaction should also be added with a No Template Control (RPA NTC), which can be further coupled with PCR (coupled NTC). PCR should also include its own NTC. It is also important to mark RPA lid strips to avoid direction inversion of lid strip. In the RPA-PCR couple, either RPA or PCR can be provided with UDG enzyme and the dTTP partially or completed replaced with dUTP. Importantly the UDG system cannot be applied to both RPA and PCR in the couple as the product of RPA is being utilized by the coupled PCR as initial template.

## Supporting information

Table S1, supportive information

## SUPPORTING INFORMATION

Table S1 provides list of the taxa utilized in laboratory evaluations of specificity and their isolate numbers, sources and origins of recovery

## ACKNOWLEDGMENT

Initial optimization of the RPA-PCR couple was accomplished during the ‘Tauti voi ei!’ project funded by Euroopan maaseudun kehittämisen maatalousrahasto (European Agricultural Fund for Rural Development/ EAFRD), Pohjois-Savon ELY-Keskus.

We specially thank the technical support team of TwistDx for introducing us with TwistAmp^®^ Liquid Basic and generously answering our questions.

