## Supplementary material for "RPA-PCR couple: an approach to expedite plant diagnostics and overcome PCR inhibitors": Table S1, supportive information

**Table S1: List of the taxa utilized in laboratory evaluations of specificity and their isolate numbers, sources and origins of recovery**

| Species | Isolate Number | Source | Country |
|---|---|---|---|
| *Phytophthora fragariae* | MRV1 | *Fragaria ananassa* | Finland |
| *Phytophthora cactorum* | PO245 | *Fragaria ananassa* | Poland |
| *Phytophthora taxon raspberry* | GE5 | Unknown (species confirmed through *atp9-nad9* sequencing) | Unknown origin |
| *Phytophthora megasperma* | GE9 | Unknown (species confirmed through *atp9-nad9* sequencing) | Unknown origin |
| *Phytophthora rosacearum* | SO18 | Unknown (species confirmed through *atp9-nad9* sequencing) | Unknown origin |
| *Phytophthora ramorum* | Ph426 | *Rhododendron catawbiense 'Grandiflorum'* | Finland |
| *Phytophthora plurivora* | Ph441 | *Rhododendron 'Marketta'* | Finland |
| *Phytophthora pini* | Ph443 | *Rhododendron 'Capistrano'* | Finland |
| *Phytophthora cambivora* | BBA 21/95-K II | *Chamaecyparis lawsoniana 'Columnaris'* | Germany |
| *Phytophthora cinnamomi* | BBA62660 | *Rhododendron* sp. | Germany |
| *Pythium sylvaticum* | ISO-VTJC | *Fragaria ananassa* | Finland |
| *Botrytis cinerea* | ISO-57C | *Fragaria ananassa* | Finland |
| *Colletotrichum acutatum* | PCF753 | *Fragaria ananassa* | Belgium |
| *Mucor hiemalis* | ISO-125-MKR | *Fragaria ananassa* | Finland |
| *Fusarium avenaceum* | ISO2-125-MKR | *Fragaria ananassa* | Finland |
| *Fusarium proliferatum* | Tiste Koe 25 | *Fragaria ananassa* | Finland |
